# Diet predicts performance trade-offs during Atlantic salmon ontogeny

**DOI:** 10.1101/2024.06.04.597289

**Authors:** Tutku Aykanat, Jan Arge Jacobsen, Kjetil Hindar

## Abstract

Understanding the mechanistic basis of life history divergence is fundamental for predictive ecology. Variation in resource use impacts life history divergence and provides a framework for studying life history trade-offs. Contrary to the theory, empirical evidence does not support that resource allocation alone mediates life history divergence in the wild, which subsequently suggests a pivotal role for resource acquisition, but this has not been empirically demonstrated in free-living populations. Here, we sampled wild Atlantic salmon (*Salmo salar*) from different age groups during their marine feeding stage and hypothesized that prey taxa (crustaceans *versus* fish) have an age-dependent effect on the condition factor. We found that salmon in their first year at sea had a higher condition factor when foraging on larger numbers of crustaceans and a lower condition factor when foraging on larger numbers of fish, and *vice versa* in older sea age groups. Provided that condition factor is a predictor of age-at-maturity, our results show that resource acquisition may be an important process shaping life history variation in Atlantic salmon via modulating age-dependent performance trade-offs. Our results also emphasize the importance of bottom-up trophic cascades for maturation age and help to better predict the maturation dynamics of wild populations in the marine environment.

## 1. INTRODUCTION

An important focus of evolutionary biology has been to identify and study the underlying mechanisms of life history trade-offs in wild populations (Reznick et al. 2000, Stearns 2000, Leroi 2001, Roff 2002, Braendle et al. 2011). By characterizing the physiological basis of the trade-offs between maturation timing and survival, the response to selection can be predicted more accurately (Roff 2002, Stearns 2011). Competitive resource allocation —the allocation of available energy to support various biological processes such as growth and reproduction—, has been suggested as the major contributor to the trade-offs dynamics in many life history models, such as in a trade-off between investing to current and future reproductive success (reviewed by Flatt et al. 2011). However, the expectation of negative phenotypic (or genetic) correlations between survival and other fitness traits lack confirmation in natural populations (Haave-Audet et al. 2022, Chang et al. 2024). Essentially, traditional trade-off models do not account for variation in resource acquisition, —the processes associated with obtaining energy from the environment to support biological processes—. Resource acquisition, however, could not only explain the observed positive phenotypic (or genetic) correlation between fitness components (Van Noordwijk & de Jong 1986, Houle 1991, Snell-Rood et al. 2015), but also provide a realistic framework to assess the mechanistic basis and the role of life history trade-offs in natural systems. For example, a recent meta-analysis in non-human animals found that differences in resource acquisition were more important than differences in resource allocation in mediating the relationship between behavior and fitness (Haave-Audet et al. 2022).

Atlantic salmon (*Salmo salar*) is a good model to study life history evolution and how life history variation is maintained (Schaffer 2003). Atlantic salmon is an anadromous species in which juveniles migrate to resourceful marine environment to enhance growth and subsequently increase reproductive success. The number of years that Atlantic salmon stay at sea prior to first reproduction (henceforth, sea-age at maturity) and fecundity are coupled because fish size and, consequently, fecundity increases with age. By extension reproductive success and mortality are inherently negatively correlated due to deceasing overall survival probability of salmon by staying longer at sea before reproduction. The maintenance of trait variation represents a classic case of fitness trade-off between reproductive success and survival: longer times at sea increase fitness via a size-dependent increase in reproductive success but at the expense of a higher risk of mortality prior to reproduction (Fleming & Einum 2010, Mobley et al. 2021). Subsequently, the species also represents a wild model to study the potential contribution of resource acquisition or allocation in mediating the optimal age structure, which also has substantial practical value for predicting the dynamics of the demographic structure of the populations. Whether resource acquisition is involved in life history trade-offs has yet to be empirically shown in wild systems.

Sea age at maturity is a threshold trait (Thorpe et al. 1998) with a strong genetic component (Barson et al. 2015, Sinclair-Waters et al. 2020), and environmental factors also play a central role in shaping the trait variation (Jonsson & Jonsson 2004, Jonsson et al. 2012). In the marine environment, Atlantic salmon achieve high growth rates and accumulate fat reserves to support energetically expensive maturation and reproduction (Jonsson & Jonsson 2011). As such, variation in resource acquisition between individuals may play a central role in shaping sea age at maturity via its effect on lipid content, which is suggested as the underlying liability trait of maturation (Rowe et al. 1991, Thorpe et al. 1998). The changes in lipid content of individuals may be mirrored by changes in condition factor, hence condition factor can predict maturation (Herbinger 1987, Herbinger & Friars 199, Duston & Saunders 1999, Peterson & Harmon 2005, Jonsson et al. 2012, Jonsson et al. 2013, Debes et al. 2021). For example, Herbinger (1987) found by studying individually marked salmon, that maturation in the autumn was associated with a high growth rate during the previous winter, and that this growth was attained by weight rather than length increase, resulting in a higher condition factor. Concordantly, it has been suggested that food web changes in the ocean may have a substantial impact on the maturation age of salmon populations (Friedland et al. 2005, Vollset et al. 2022), and salmon populations may show evolutionary responses to changes in food availability (Czorlich et al. 2022). At sea, the diet of sub-adult Atlantic salmon (the marine stage prior to maturation) mainly consists of crustaceans and fish, with large spatiotemporal variation (Jacobsen & Hansen 2001, Rikardsen & Dempson 2010). These two major taxonomic groups have fundamentally different properties as potential prey, such as size, behavior, and availability, which ultimately require different foraging strategies to utilize them (Mittelbach & Persson 1998, Ferry-Graham et al. 2002, Sanchez-Hernandez et al. 2019). Feeding also has an ontogenetic component: crustaceans-based diet in early marine phases switches to a more piscivorous diet as individuals grow larger. However, there is a considerable overlap in the distribution of prey groups between different sea ages (Jacobsen & Hansen 2001).

Therefore, age-dependent dynamics of diet intake and how it affects condition factor, a strong predictor of maturation timing (Herbinger & Friars 1991, Jonsson et al. 2012), is unclear in wild Atlantic salmon.

The ontogenetic diet shift is an adaptive trait in which an optimal feeding strategy is attained to supply the increasing energetic demands of a larger body, and is mostly linked to changes in habitat and life-stage transitions (Villéger et al. 2017, Sanchez-Hernandez et al. 2019). However, few studies have quantified inter-individual variation in ontogenetic diet dynamics (Mittelbach & Persson 1998, Sanchez-Hernandez et al. 2019), with no assessment of potential performance trade-offs in free-living populations. An ontogenetic diet shift in the marine environment implies that the net adaptive value of energetic gain obtained from a particular feeding regime (e.g., diet type) changes as a function of ontogeny. As such, acquiring optimal resource type increases the net energy gain, which may facilitate lipid accumulation and condition factor (Jonsson et al. 2013), and subsequently probability of maturing (Herbinger & Friars 1991, Rowe et al. 1991, Thorpe et al. 1998). Therefore, resource acquisition may be suggested as a functional ecological process explaining the dynamics of age at maturity, by demonstrating an age dependent relation between performance (condition factor) and diet composition (Haave-Audet et al. 2022, Chang et al. 2024).

**Fig. 1.**
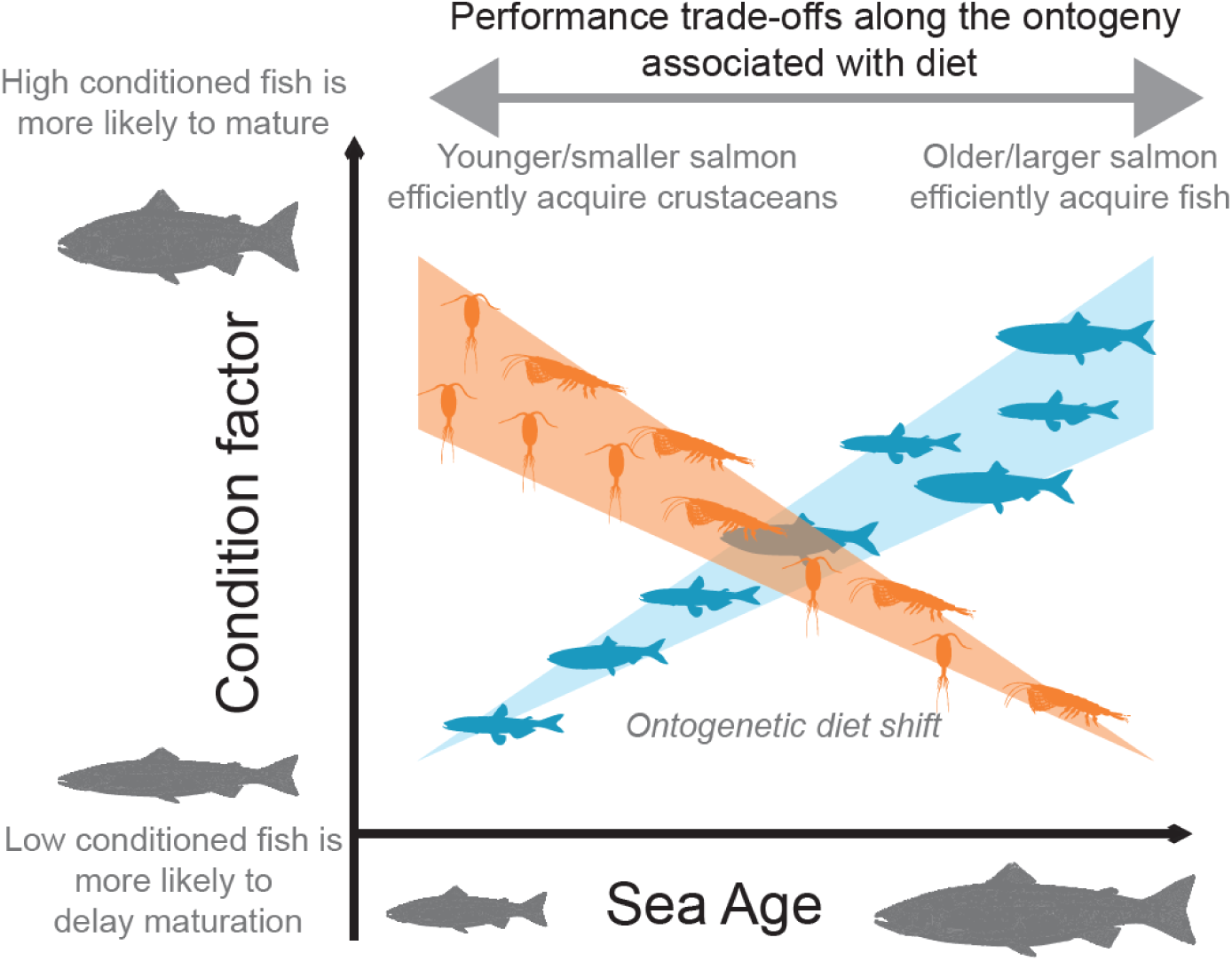
Conceptual model that links performance (condition factor) to diet type (crustacean versus fish) of sub-adult Atlantic salmon during the marine feeding stage.

Based on the above predictions, we hypothesized there is an ontogeny-dependent performance trade-off for consuming different diet items (Fig. 1). Specifically, we hypothesize that eating smaller food items early in the marine stage is an optimal feeding strategy for smaller-sized salmon, and increases their probability of maturation in the following year by facilitating lipid depositions, which may be predicted by the condition factor (Herbinger & Friars 1991, Todd et al. 2008). Lipids are essential, for example, during vitellogenesis (egg production), which comprises a substantial amount of energy expenditure of females during the reproduction period (Jonsson & Jonsson 2003), as well as for other energetically expensive migration and reproductive activities (Jonsson & Jonsson 2011). Concordantly, lipid content has been suggested as the underlying, liability trait that triggers the developmental switch to initiate the maturation the following year (Thorpe et al. 1998). Likewise, we hypothesize that switching to larger food items is more beneficial to older, larger-sized salmon, leading to higher net energy gain, also to support future reproduction activities. Such age dependent trade-off between performance (condition factor) and diet composition would provide an ecological explanation of how a life history trait (age at maturity) is mediated by resource acquisition variation (marine diet composition). As such, we quantified the diet of wild Atlantic salmon sub-adults sampled in the oceanic feeding ground, quantified their stomach contents (consisting of crustaceans and fishes, Jacobsen and Hansen 2001), and compared the age-dependent changes in condition factor in relation to diet.

## 2. MATERIALS & METHODS

### 2.1. Study system

Atlantic salmon used in this study were originally described by (Jacobsen & Hansen 2001), and the final data were identical to Aykanat et al. (2024) except that we retained N=68 individuals that were filtered out in the prior study due to missing genotype information in two focal life history loci. Briefly, the dataset comprises 1604 individuals collected from the Northeast Atlantic Ocean in their ocean feeding ground North of the Faroe Islands, of which 178, 995, and 431 are from 1SW, 2SW, and 3SW sea-age groups, respectively (SW, “sea-winter” is the number of winters a fish has been in the marine environment and is equivalent to the sea age). Samples were collected at 55 locations during fishing periods in autumn (November and December) and winter (February and March) in 1992-93 and 1993-94 (Fig. S1). The fish were genetically assigned to large-scale reporting groups (Ireland and the UK, Southern Norway, and Northern Norway; the latter encompassing fish assigned to Barents Sea drainages (O’Sullivan et al. 2022), which descibes broad phylogenetic groups associated with large geographical regions. Compared to the original study (Jacobsen & Hansen 2001), which comprised of N=1765 wild individuals from the 1992-93, and 1993-94 fishing periods, we excluded 161 individuals, of which N=34 did not have any material to genotype (i.e. no scales), N=9 did not have sex information, N=15 had sea age larger than 3SW, and 103 were not assigned to the reporting groups indicated above (out of 103, N=88 was assigned to North American phylogenetic groups). Individual salmon had their fork lengths measured in centimeters, and gutted, and gutted fish were weighed in kilograms to two decimal places. The gut contents were identified to taxon level and enumerated. Fish scales were used to determine sea age (via scale reading), and for DNA extraction and genotyping. The fish sampled in this study were sub-adults with no phenotypic indications of maturation. An independent, earlier study from the Faroese feeding grounds indicates that 80% of individuals are likely to mature in the following autumn, but these estimates are based on limited number of tag recapture studies, or based on assessing the maturation probability via hormonal expression using hatchery reared salmon as the baseline (Youngson & McLay 1984), hence may not be reliable.

### 2.2. Data analysis

We employed a two-step multi-model inference approach using linear mixed models, using condition factor as the response variable. Condition factor is a strong predictor of lipid content in Atlantic salmon (Herbinger & Friars 1991, Todd et al. 2008) which is suggested as the liability trait of the maturation timing (Thorpe et al. 1998). In this framework, we first identify the most parsimonious null model structure by comparing the fit of the models parameterized with different combination of non-focal variables, and their interactions. The global null model structure is as follow:

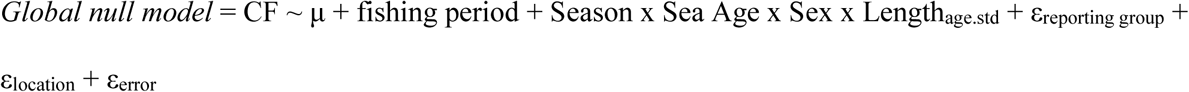

Whereby, μ is the intercept and εerror is the residual errors. The condition factor (CF) is Fulton’s condition factor K, (K=10^5^ w / l^3^), in which w is the gutted weight in kilograms divided by 0.9, to account for gut weight, and l is fork length in cm (Fig. S2). Fulton’s condition factor is a common condition index that has been a sufficient standard in physiological studies in fishes (Stevenson & Woods 2006). The condition factor is based on the weight obtained from gutted fish; therefore, it reflects the condition factor associated with body weight excluding the gut weight and visceral fat and gonadal tissue weights. This is pertinent since subcutaneous adipose tissue and musculature lipids have been suggested as the main energy reserves during the spawning period (Jonsson et al. 1997).

The model included season, fishing period, sea age, sex, and age-standardized length (Lengthage.std) as fixed effects. Spatial variation in sampling effort was accounted for by including sampling location as a random effect (εlocation, N=55). Likewise, variation due to geographical origin of individuals was accounted for by including reporting groups (εreporting group, N=3) as random terms. Lengthage.std was scaled to have a mean of zero and a standard deviation of one to improve convergence. All four-way combinations of season, sex, sea age, and age-standardized length and their reduced models were fitted and compared. Models were fitted by parameterizing sea-age both as a continuous or factorial variable (Table 1). All models were then compared using corrected Akaike Information Criteria (AICc), which corrects for bias for small sample sizes. The model with lowest AICc value is considered as the most parsimonious null model and used in the following analyses.

**Table 1:**
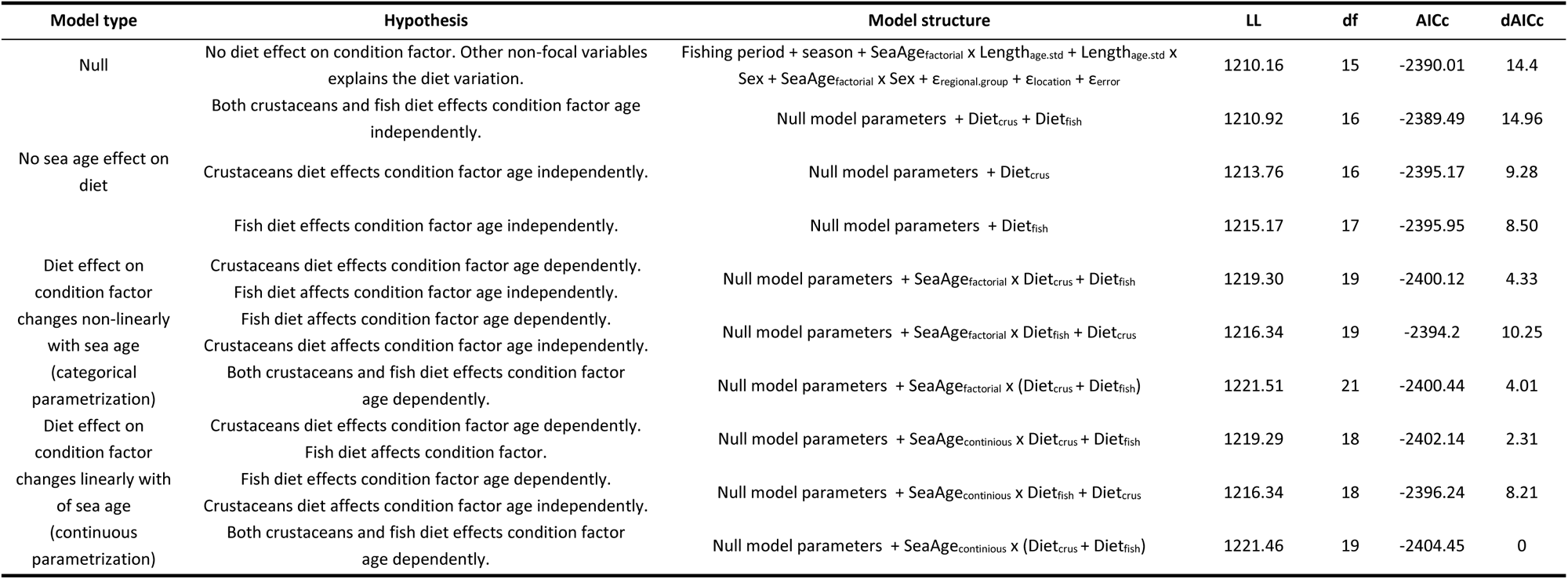
Parsimony of models with different parametrizations. Null model is the most parsimonious structure containing non-focal terms, and this structure included in hypothesis-specific models. LL indicates log likelihood of the model.

Next, on top of the most parsimonious null model, we fitted models that also include combinations of focal variables (Dietcrus and Dietfish) and their interaction with sea age, the fit of which was then compared using AICc. These models reflected hypotheses explaining the diet type effect on condition (Table 1). The global model structure is as follow:

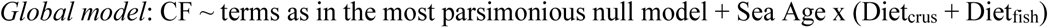

Whereby, Dietcrus and Dietfish are log-normalized number of diet items in the stomach. Both crustacean and fish diets had highly left-skewed distributions with many zero entries (i.e., empty stomach, Fig. S3). To avoid undefined zero values after log transformation, we added 1 to the number of diet variables. As above, sea-age was modelled both as a continuous or categorical variables.

All models were run using the glmmTMB package (version 1.1.5, Brooks et al. 2017) in R version 4.1.0 (R Core Team 2019), using a Gaussian error structure. Null model selection and AICc calculations was performed using MuMin package (version 1.48.4, Barton et al 2024). Marginal means for genotype effects and their confidence intervals were calculated using the emmeans package, and significance was assessed by t-test (version 1.8.5, Lenth, 2023). Model diagnostics were performed using the DHARMa package in R (version 0.4.6, Hartig 2022).

### 2.3. Post-hoc analyses

We implemented a number of post-hoc analysis in which we revise original parameterization of terms to further break down the structure of the diet effect on condition.

#### 2.3.1. The effect of different taxonomic groups within crustaceans

We evaluated whether any taxonomic groups within crustaceans drive the relationship associated with the crustaceans diet, by partitioning crustaceans diet into lower taxonomic groups, provided that the numbers within a specific taxon were sufficiently large. As such, we implemented the most parsimonious model structure, but by replacing Dietcrus term with either DietHyperiid.amphipods or DietEuphausiids, which are the log-normalized number of hyperiid amphipods (*Themisto* spp.) and euphausiids in the diet, respectively. These two taxa comprise the majority of the crustaceans in the stomach (Table S1), and were substantially different in size, with the latter being larger than the former (Table S1). We did not implement the same approach in fishes due to small number of prey items per species (Table S1).

#### 2.3.2. Partitioning the underlying resource acquisition processes

The relationship between the condition factor and number of prey items might be driven by different ecological processes associated with foraging, such as active foraging frequency or foraging efficiency (Aykanat et al. 2024). We assessed the contribution of these two possibilities by repeating the analysis after restructuring diet variables to account for these factors. As such, the presence or absence of prey was used as a proxy for active foraging frequency, using the model structure described above, but replacing the diet variable with a Boolean parameter, for crustaceans and fish in two separate binomial models. Using the data with which diet was present, log-transformed number of diet items was modelled as a proxy for foraging efficiency (when actively foraging for prey), for crustaceans and fish in two separate models.

#### 2.3.3. Comparing the importance of weight vs number of diet in the stomach

The number and total weight of the diet items in the stomach were highly correlated (Fig. S4); thus, any effect due to the number of diet items could be attributed to diet weight. To assess whether the number or total mass of the diet better explain the change in the condition factor, we implemented the same model as above but used total diet weight as a factor instead of the number of prey items acquired.

## 3. RESULTS

### 3.1. Null model structure and parametrization of sea age

The most parsimonious null model included fishing period, season, lengthage.std, sex, and sea age as parameters, and second order interactions between lengthage.std, sex, and sea age, (Table S2). Notably, a model that parameterizes sea age categorically was substantially better than a model with continuous parametrization (ΔAICc = 67.18, Table S3). However, we modelled sea age continuously when interacting with diet variables. Such a model was more parsimonious than a model that parameterizes sea age continuously (ΔAICc = 87.78) or categorically (ΔAICc = 4.01) across all the interactions. In all three models, main results regarding age-dependent diet association on condition factor was qualitatively similar (data not shown).

**Fig. 2.**
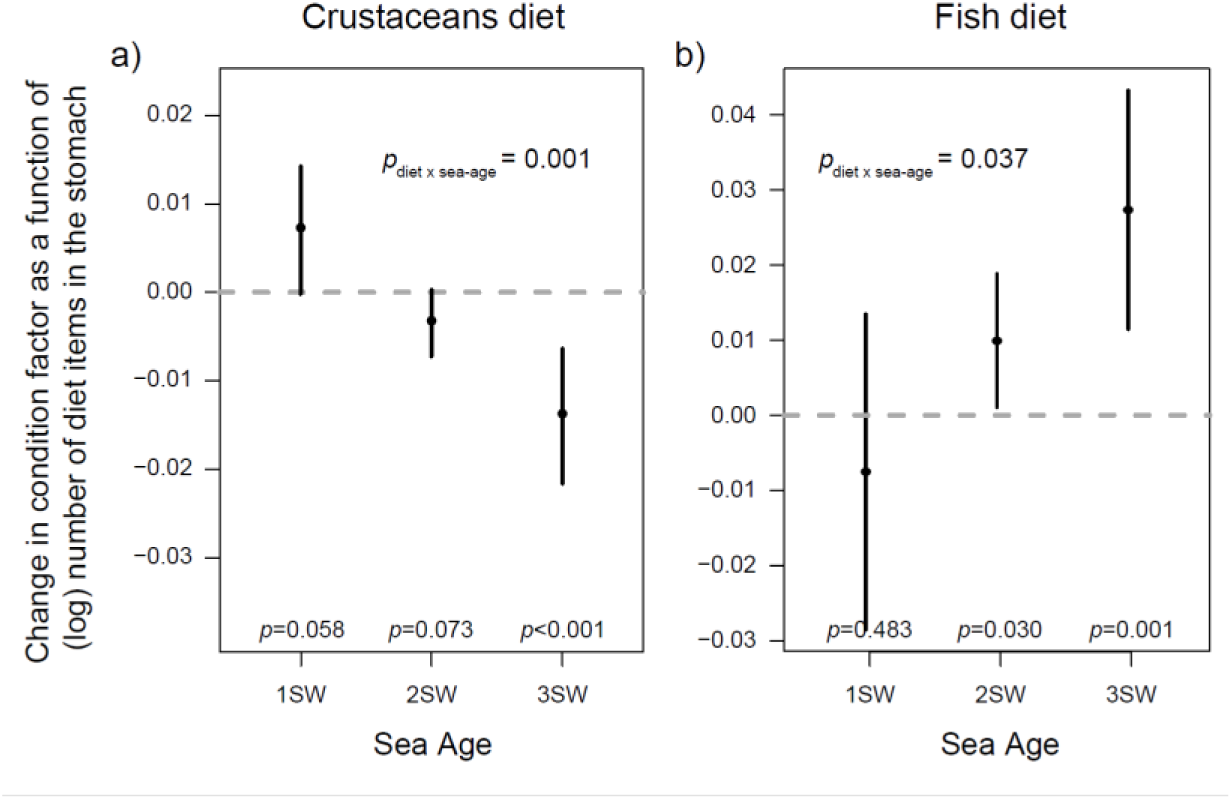
Marginal effect of the number of crustaceans (a) and fish (b) prey items (log scale) on condition factor in Atlantic salmon sub-adults in the marine environment. Dashed line indicates a zero slope, no relation between the number of diet items and condition factor. Error bars indicate 95% confidence intervals.

### 3.2. Age dependent diet effect on condition factor

Our analysis supported the hypothesis that suggests an age dependent diet effect on condition factor (Table 1). In this model, the number of diet items in the stomach had a significant age dependent association with condition factor, but the direction of the age effect was in contrasting for crustaceans and fish diets. In other words, the age dependent slope of the relation was negative for crustaceans diet (for crustaceans diet by sea age interaction: t-test, *t1585* = -3.208, *p =* 0.001, Fig. 1a), but it was positive for fish diet (for fish diet by sea age interaction *t1585* = 2.088, *p =* 0.037, Fig. 1b). Within 1SW age group, the number of crustaceans in the diet was positively associated with condition factor (*t1585 =* 1.901, *p =* 0.058), but the relation was increasingly negative as the age of salmon progresses (for 2SW: *t1585 =* -1.794, *p =* 0.073, for 3SW: *t1585 =* -3.572, *p <* 0.001. Fig. 2a). The number of fish in the diet did not affect the condition factor in the 1SW age group (*t1585 =* -0.702, *p =* 0.48), but the relation was increasingly positive as the age of Atlantic salmon progresses (for 2SW: *t1585 =* 2.177, *p =* 0.030, for 3SW: *t1585 =* 3.367, *p <* 0.001, Fig. 2b, Fig. S5). The scenario where crustaceans and fish diets were modelled age dependently with continuous parameterization of sea age significantly outperformed any other alternative hypotheses (Table 1).

### 3.3. Age dependent sex and length effects on condition factor

Sex and length also exhibited age-dependent patterns, which had an apparent non-linear dependency with sea age (Fig. 3). Notably, there was a substantial reversal of sex-dependent changes in the condition factor between the 1SW and 2SW age groups (Fig. 3a). Males had significantly higher condition factors in the 1SW age group (*t1585 =* 2.693, *p =* 0.007) and significantly lower condition factors in the 2SW age group (*t1585 =* -5.254, *p <* 0.001), and there was no difference between sexes in the 3SW age group (*t1585 =* 0.026, *p =* 0.98, Fig. 3a). Length was positively associated with the condition factor positively in the 2SW age group (*t1585 =* 5.040, *p* <0.001) but the association was not significant in other age groups (Fig. 2b). As expected, winter was associated with a lower condition factor (*t1585 =* -8.123, *p <* 0.001), and there was a marginal fishing-period effect (*1585 =* -1.989, *p <* 0.047, see also Table S2).

**Fig. 3.**
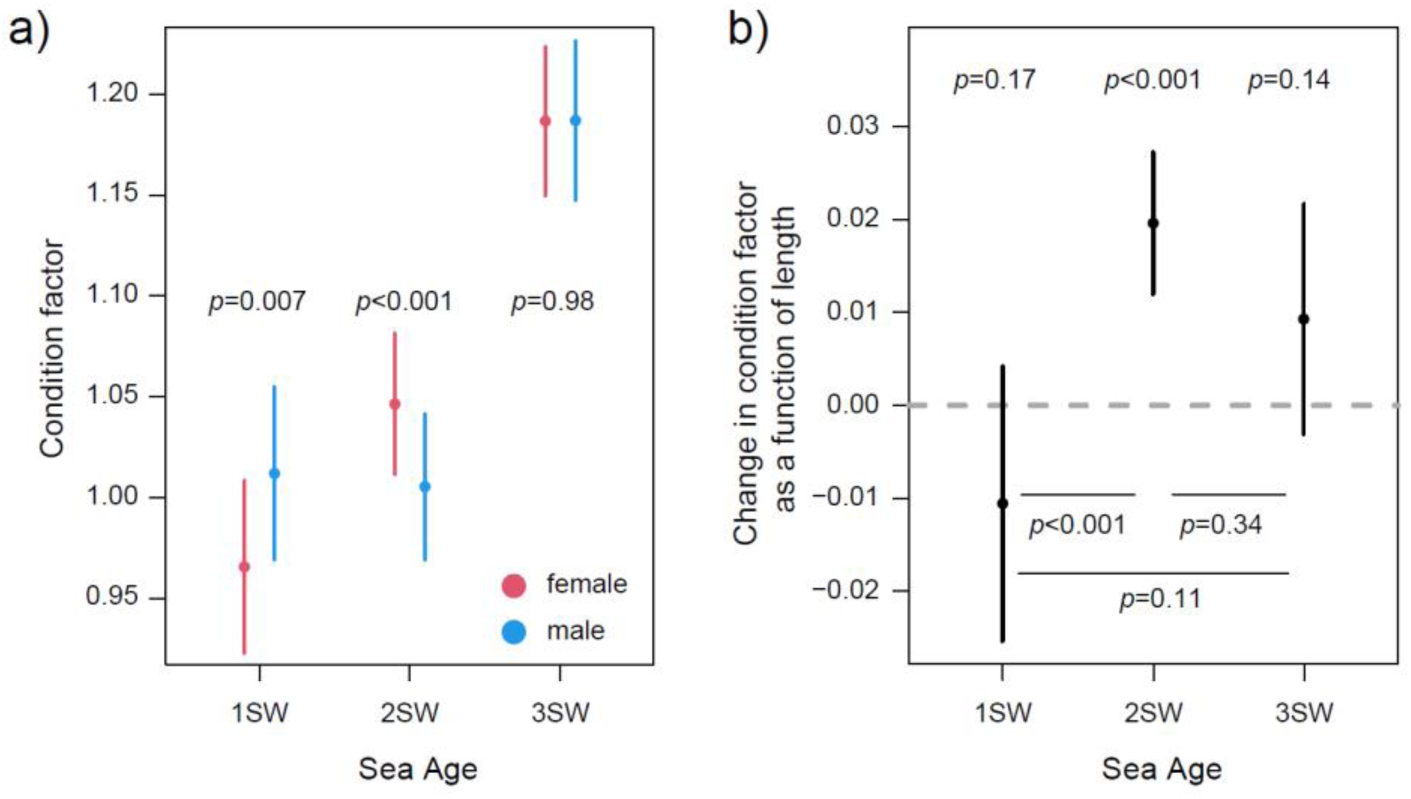
Marginal effects of sex (a) and length (b) on condition factor in Atlantic salmon sub-adults in the marine environment. Dashed line in b indicates a zero slope, no relation between (age standardized) length of salmon and condition factor. Error bars indicate 95% confidence intervals.

### 3.4. Taxon specificity of crustacean diet effect

The crustacean diet in the stomach consisted of a few major taxonomic groups, the majority of which were hyperiid amphipods (∼93% and 69% by number and weight, respectively, and mostly *Themisto* species) and euphausiids (krill, ∼ 6% and 15% by number and weight, respectively, Table S1). When the number of diet specific to these taxa was separately parameterized in the model, instead of crustaceans as a whole, hyperiid amphipods, but not krill numbers, explained the ontogenetic diet effect on the condition factor (Fig. 4, Table S4), suggesting that the overall pattern was driven by hyperiid amphipods acquisition. As such, a model that contained hyperiid amphipods explained the data substantially better than the original model that had parameterized crustaceans as a whole (ΔAICc = 2.53, Table S4). Concordantly, the age-dependent slope of the association between the number of hyperiid amphipods in the diet and condition factor was strongly significant (for hyperiid amphipods diet by sea age interaction: *t1583 =* -3.621, *p <* 0.001). The condition factor of 1SW was significantly higher when the diet contained more hyperiid amphipods (*t1583* = 2.146, *p =* 0.032), but the relation was directly opposite direction in 2SW (*t1583 =* -2.112, *p =* 0.035) and 3SW age groups (*t1583 =* -3.941, *p <* 0.001, Fig. 4a). A parallel analysis for the fish diet was not performed because the number of diet items was prohibitively small when the fish diet data were partitioned into smaller taxonomic groups.

**Fig. 4.**
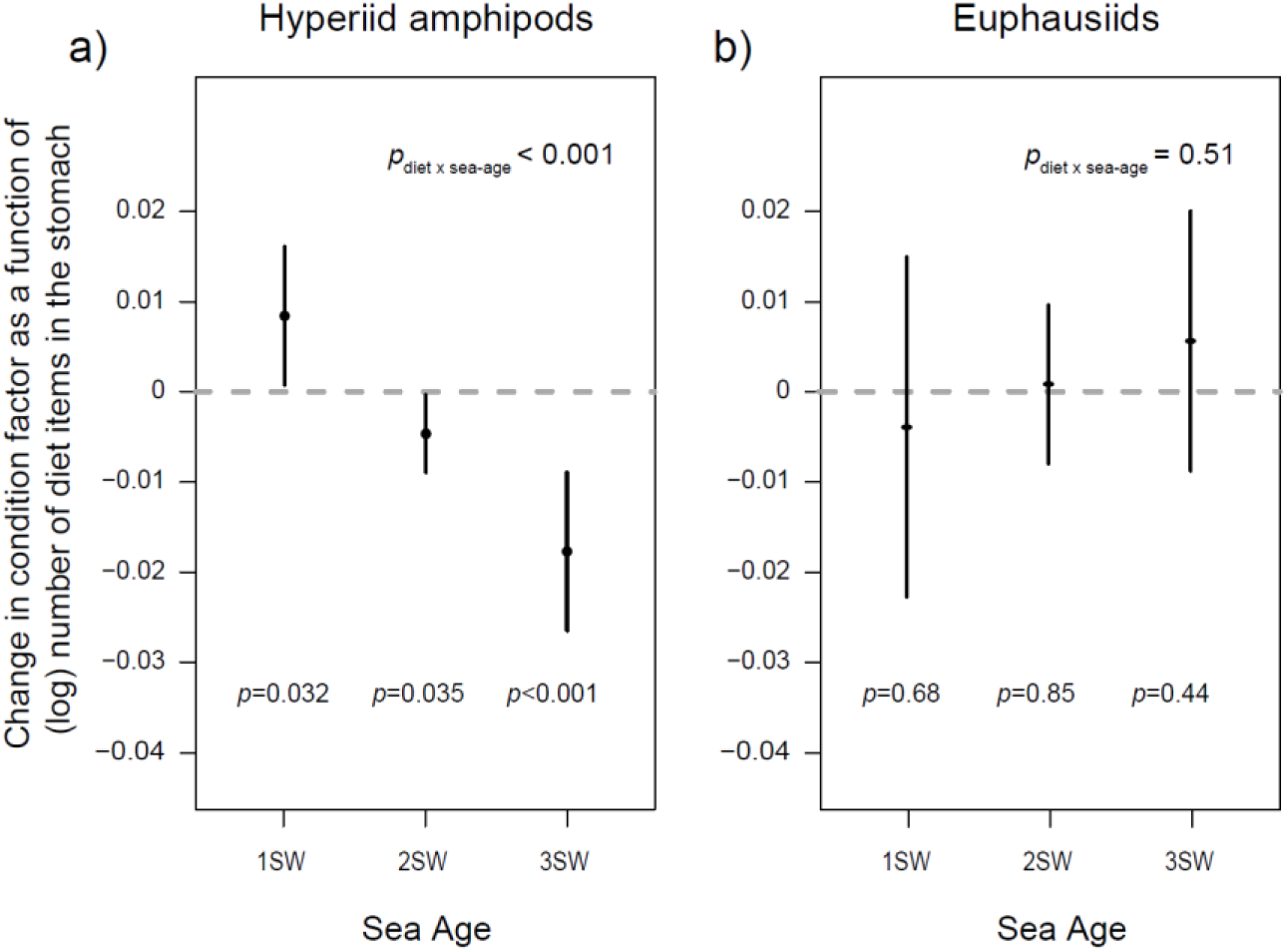
Marginal effects of the number Hyperiid amphipods (a) and Euphausiids (b) prey items (log scale) on condition factor in Atlantic salmon sub-adults in the marine environment. Dashed line indicates a zero slope, no relation between the number of diet items and condition factor. Error bars indicate 95% confidence intervals.

### 3.5. Contribution of diet weight vs. number of diet items on condition factor

The relationship between the condition factor and number of prey items might be driven by different ecological processes associated with foraging, such as foraging frequency or foraging efficiency (Aykanat et al. 2024). We assessed the contribution of these two processes after restructuring the response variable into two parts, first to quantify foraging frequency (approximated with a Boolean diet presence and absence), and second to quantify foraging efficiency (i.e. approximated with log-transformed number of diet items, when diet was present in the stomach), and by repeating the analysis above. The effects of diet on the condition factor were qualitatively similar to each other and to the original models for both crustaceans and dish diets (Fig. S7), ruling out the sole contribution of any of the two processes.

When the total diet weight was parameterized in the model, instead of the number of prey items acquired, it did not explain ontogenetic effects in the diet (Fig. S6), suggesting that number of diet items but not weight were the primary factor associated with the ontogenetic effect.

## 4. DISCUSSION

We demonstrated that resource acquisition predicts ontogenetic performance trade-offs via its effect on condition factor. Competitive resource allocation has been the major mechanism suggested to mediate life history trade-offs within species but lacks evidence in natural systems (Flatt & Heyland 2011, Haave-Audet et al. 2022, Chang et al. 2024). The theoretical premise of resource acquisition in shaping life history trade-offs dynamics has been acknowledged directly (Flatt & Heyland 2011), or indirectly across different approaches, as a potential outcome through the effect of signaling networks (Stearns 2011) or via the effect of changing environmental plasticity to couple-decouple the correlations between life history traits. However, the empirical studies involve only common garden designs under controlled conditions (Messina & Fry 2003, Ernande et al. 2004, Lailvaux et al. 2005). Our results, obtained in the wild, demonstrate a link between the numbers of different prey species in the diet and condition factor, a strong predictor of the liability trait of age-at-maturity. Therefore, we suggest that Atlantic salmon life history dynamics may be shaped by resource acquisition dynamics in the wild via the availability and composition of different prey species.

Among crustacean taxa, hyperiid amphipods (mostly *Themisto* spp.), but not euphausiids (krill), were associated with an age-dependent decrease in condition factor. Hyperiid amphipods are much more abundant prey type in the marine feeding grounds of the Northeast Atlantic (Jacobsen & Hansen 2001) but are smaller than krill or other crustaceans taxa observed in the diet (Table S1). Optimal foraging theory predicts that organisms select prey that maximize net energy gain (Pyke et al. 1977). It is plausible that acquiring hyperiid amphipods becomes increasingly energetically expensive as salmon get older and larger, perhaps due to size dependent changes in morphology and swimming ability (Sanchez-Hernandez et al. 2019). In contrast, the influence of fish diet on the condition factor increases with sea age. Consuming fish increases the condition factor only in older sea age groups, suggesting that some energetic constraints associated with smaller fish size exist, which may be associated with gape size (Keeley & Grant 1997, Scharf et al. 2000), or size dependent attributes that makes younger fish to foraging for fish less efficient (Webb et al. 1984, Sanchez-Hernandez et al. 2019, see also below). Taken together, foraging predominantly on hyperiid amphipods at younger ages and shifting to foraging fish prey at older ages may signify an age-dependent optimal foraging strategy for Atlantic salmon, at least in the environmental settings that the sampling have taken place.

Combined with the fact that winter condition factor at sea is a strong predictor of maturation timing in the following season (Herbinger 1987, Herbinger & Friars 1991), our results may have significant implication for understanding the maturation dynamics of Atlantic salmon. For example, we predict that a younger age structure at maturity will be associated with relatively higher crustacean abundances (mainly hyperiid amphipods). Likewise, relatively higher fish prey abundances might negatively affect condition factor at younger age groups, leading to delayed maturation and older age structure at maturity. Vollset et al. (2022) suggested that broad-scale changes in oceanic conditions led to an ecological regime shift that subsequently affected the age at maturity. In their 28-year time series data, the decrease in planktonic biomass was highly correlated to an increase in the proportion of older sea age groups (Vollset et al. 2022), which corroborates our prediction. In their study, there was a strong covariation across several abiotic and biotic factors during the regime shift, making it challenging to break down the drivers of this change further. Our study linked individual-based diet patterns to condition factor across a heterogeneous spatial distribution of Atlantic salmon and its prey, and within a narrow temporal window, thus providing an assessment that is independent from the effect of temporally covarying factors, such as temperature. As such, we can suggest, among other factors outlined in Vollset et al. (2022), that decreased crustacean abundances are likely the driver of observed older age structure.

While the ontogenetic diet shift towards a more piscivorous diet has been demonstrated in this system earlier (Jacobsen & Hansen 2001), its effect on condition factor was not known until this study. Based on our results, one can suggest that the net benefit of a crustaceans oriented diet is diminishing with age, while a fish-based diet becomes more rewarding. Since the ontogenetic relationship is driven by prey numbers, but not mass (Fig. S6), we can postulate that the age-dependent diet benefits are caused not by post-acquisition processes, such as differences in prey energy content or potential differences in nutrient stoichiometry between diet types, but by differences in energy use prior to or during prey acquisition. The result that this relationship is mostly driven by the smaller (and more numerous) hyperiid amphipods (Fig. 4) further supports this notion. Therefore, our results are consistent with a scenario that foraging for smaller crustaceans becomes increasingly energetically unsustainable for larger salmon. Similarly, we suggest that foraging for fish is only energetically advantageous for larger salmon, which may be due to size-dependent changes in morphology or behavior (Smith & Skúlason 1996). For example, gape size may restrict smaller salmon from capturing larger fish (Keeley & Grant 1997, Scharf et al. 2000), or they may not sustain cruising speeds that are required for foraging for fish prey as efficiently as larger salmon (Webb et al. 1984, Sanchez-Hernandez et al. 2019).

The ontogenetic effect of diet on condition was not solely explained by age-dependent changes in foraging strategy, as demonstrated by similar effects between when diet was parameterized as presence/absence (as an indication of frequency of active foraging), or total prey numbers in the diet (when diet items were non-zero, and indication of foraging outcome). This suggests that ontogenetic effects are independent of the foraging strategy. Interestingly, Aykanat et al. (2024) suggested that the *vgll3* locus, a major locus influencing age at maturity, was associated with age-dependent differences in crustaceans foraging frequency, but not foraging outcome. Fish with *vgll3* early maturing genotype appeared to gain similar amounts of food with less foraging activity, suggesting that an increase in acquisition efficiency may be one of the factors that allow salmon with the *vgll3* early genotype to mature earlier, that is, by allocating surplus resources towards reproductive investment (Aykanat et al. 2024). However, they did not find any evidence of an age-dependent association between condition factor and *vgll3* genetic variation. These results suggest that the association between diet and condition factor in the present study likely has an independent physiological pathway than how *vgll3* locus effects diet.

Similar to diet, our results show that sex has an ontogenetic effect on condition factor, in which 1SW salmon males have a higher condition factor, whereas this pattern reversed for 2SW age group (Fig. 2b). In Atlantic salmon, males mature more often as 1SW, and females more often delay maturation to achieve a larger body size since size is a stronger determinant of reproduction success in females via its effect on fecundity (Fleming & Einum 2010). Similarly, energy expenditure during spawning is higher among younger males than females but increases more by size in females than in males (Jonsson & Jonsson 2003, Persson & De Roos 2006). The reversal in sex-specific differences in condition factor between 1SW and 2SW age groups (Fig. 4) is consistent with the indicated maturation trends above, providing that condition factor predicts maturation via its effect on the lipid conetnt. This result is also consistent with the proposed sex-specific variation in the liability distribution of of age at maturity, which explains the emergence of sex-specific dominance reversal (Reid 2022), an adaptation to offset sex differences in optimal age at maturity in Atlantic salmon (Jonsson & Jonsson 2003, Fleming & Einum 2010, Barson et al. 2015).

Linking stomach content to general resource acquisition patterns has sometimes been criticized because it is only a snapshot of the diet, a measure of a very recent feeding activity, as opposed to methods such as stable isotopes that can assess long-term trends (e.g., Power et al. 2023). While integration of multiple approaches (that includes analysis of stomach content, and stable isotope and fatty acid profiling) may help to better understand complex trophic relationships (Nielsen et al. 2017), a careful formulation of hypotheses and statistical methods could avoid limitations associated with sole use of stomach content data (Nielsen et al. 2017, Amundsen & Sánchez-Hernández 2019). As such, our results remarkably explain an ontogenetic diet shift in the species (Klemetsen et al. 2003), and provide a well-supported and predictable relationship with condition factor, a body index that should not be affected by only very recent feed intake. Hence, our results confirmed that stomach content analysis reflects general feeding patterns.

The sampling site for this study was one of the major marine feeding grounds for European salmon populations, especially for older, 2SW and 3SW sea age groups (Jacobsen & Hansen 2001, O’Sullivan et al. 2022). However, many salmon populations from geographically distant localities, such as North America and the Baltic Sea, utilize other marine feeding areas, and salmon of the youngest sea age groups (1SW) may inhabit marine feeding areas that are closer to their native rivers (Rikardsen & Dempson 2010, Gilbey et al. 2021, Rikardsen et al. 2021). Subsequently, prey compositions may be substantially different in those habitats which potentially different factors effecting feeding and maturation dynamics. Therefore, the age-dependent diet performance relation that we observed here, and its potential effect on maturation may not readily be generalized across species range.

Data collected in the wild has high dimensionality. Our model contains several parameters that could otherwise be controlled under common garden conditions. While important, testing all possible interactions between covariates (which may covary with focal covariates) is not feasible owing to the number of possible combinations, and unobserved variables are also likely important for diet variation. Nonetheless, our model appears robust to the exclusion of observed variables from the model (see Results). Furthermore, despite a strong statistical support, and multiple lines of indirect evidence supporting the main conclusion, such as observing an ontogenetic diet shift that is common in salmonids (Klemetsen et al. 2003) and functional morphological variation associated with size and diet type, the age-dependent diet effect on condition factor is based on correlation of observational data.

Ideally, controlled experiments are required to confirm the causality, but this may be unfeasible for this case, given impracticalities, at the very least, associated with rearing salmon long periods and using forage species as food sources.

## 5. CONCLUSIONS

In this study, we provide a clear case of an ontogenetic performance trade-off associated with diet in a freely roaming species across large geographical areas. The empirical evidence here supports the notion that trade-offs mediating life history divergence could be explained by variation in resource acquisition in the wild, and could explain the lack of genetic or phenotypic correlations observed between life history traits. Our study provides a mechanistic basis to better predict Atlantic salmon maturation dynamics and performance trajectories across large spatiotemporal scales. Climate-induced effects on Atlantic salmon performance and demography have been suggested previously (Jonsson & Jonsson 2004, Todd et al. 2008, Friedland et al. 2009, Vollset et al. 2022). However, covariation between critical biotic and abiotic physiological factors (such as temperature and forage composition and abundances) in longitudinal data is insufficient to identify primary factors driving these patterns (Friedland et al. 2009). Our results contribute to the knowledge by providing a spatial analysis in which the effect of forage species (diet) on performance (condition factor) could be factored out from covarying factors that intrinsically exist in longitudinal data structures. Our results indicate bottom-up trophic dynamics as likely underlying factor that links large-scale ecological regime shifts to changes in maturation patterns (Vollset et al. 2022). Our study also highlights the importance of exploiting and repurposing old but gold datasets to address novel ecological questions.

## Acknowledgements

This study was funded by the Research Council of Finland (grant nos. 328860, 353388, and 325964 to TA) and the Research Council of Norway (project number 280308 to KH). We thank three anonymous reviewers whose comments greatly improved the quality of the manuscript.

## Author contributions

TA conceptualized the paper. KH organized sample retrieval from scale archives at NINA. JAJ coordinated the Faroese fieldwork and collected diet data. TA conducted the statistical analyses. TA and KH drafted the MS and all coauthors contributed to subsequent drafts.

