## Supplementary Materials for "Diet predicts performance trade-offs during Atlantic salmon ontogeny"

for

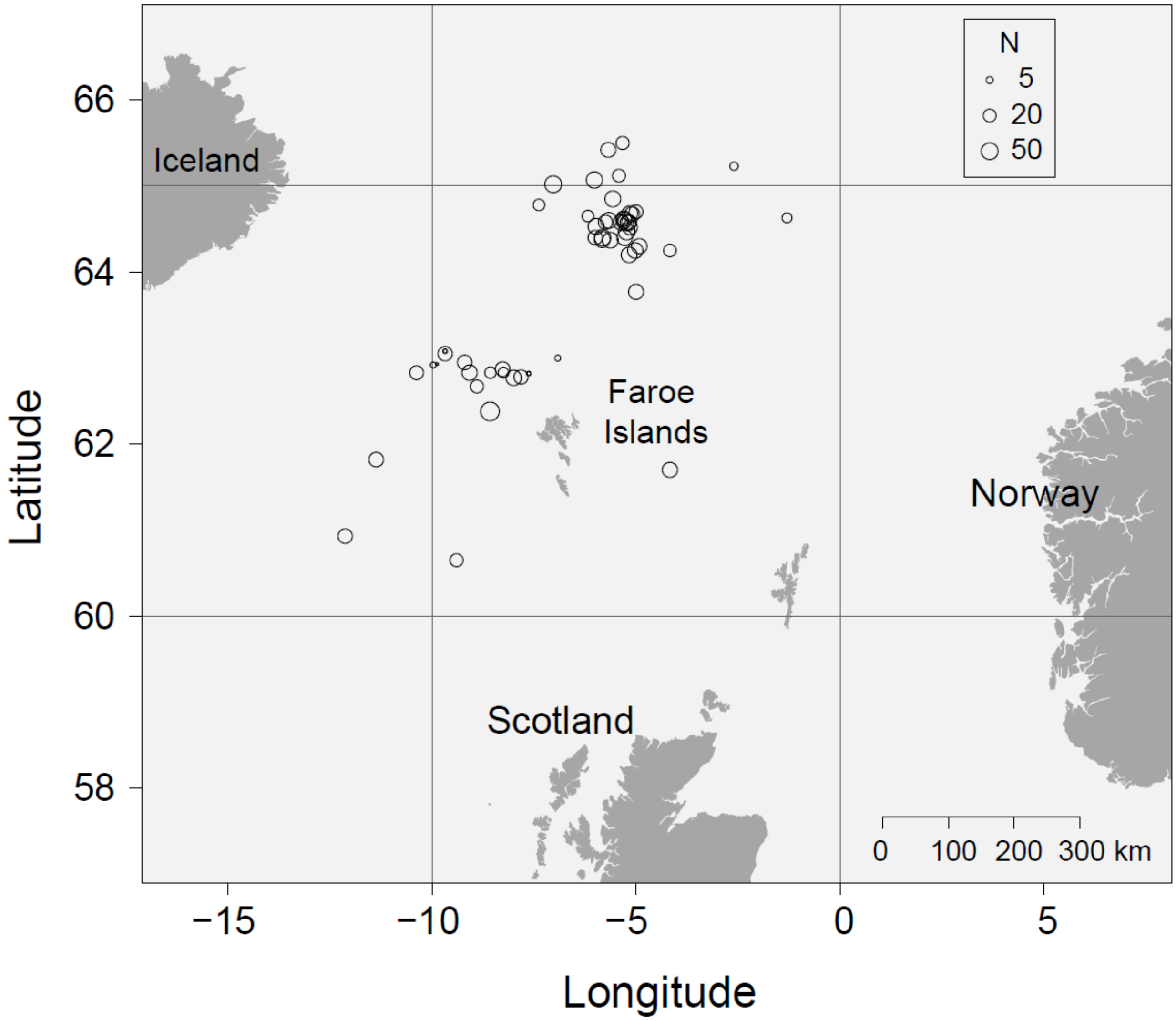


**Fig. S1.** Sampling locations (N=55) and sample sizes (N=1604) used in this study.

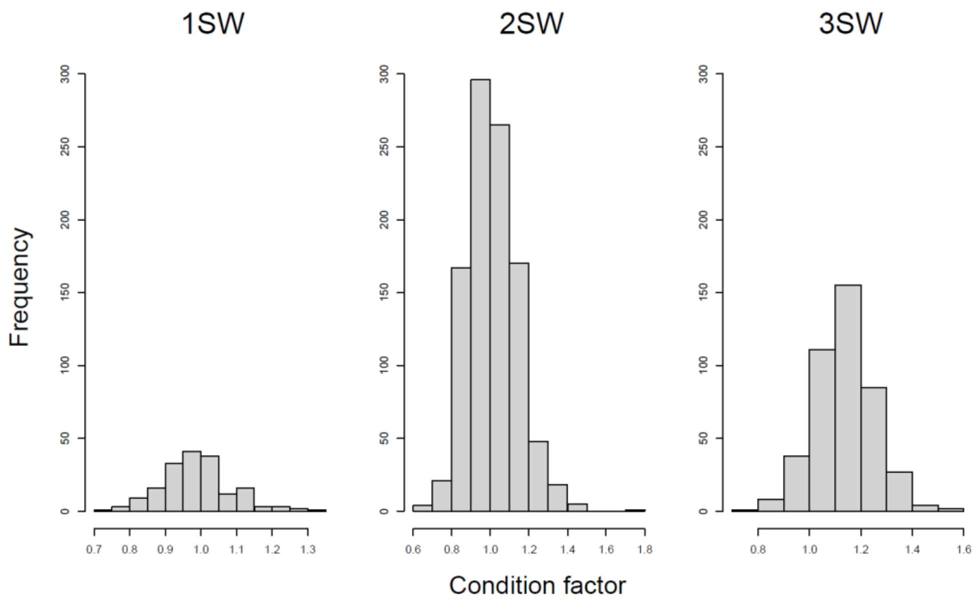


**Fig. S2.** The distribution of condition factor across sea-age groups.

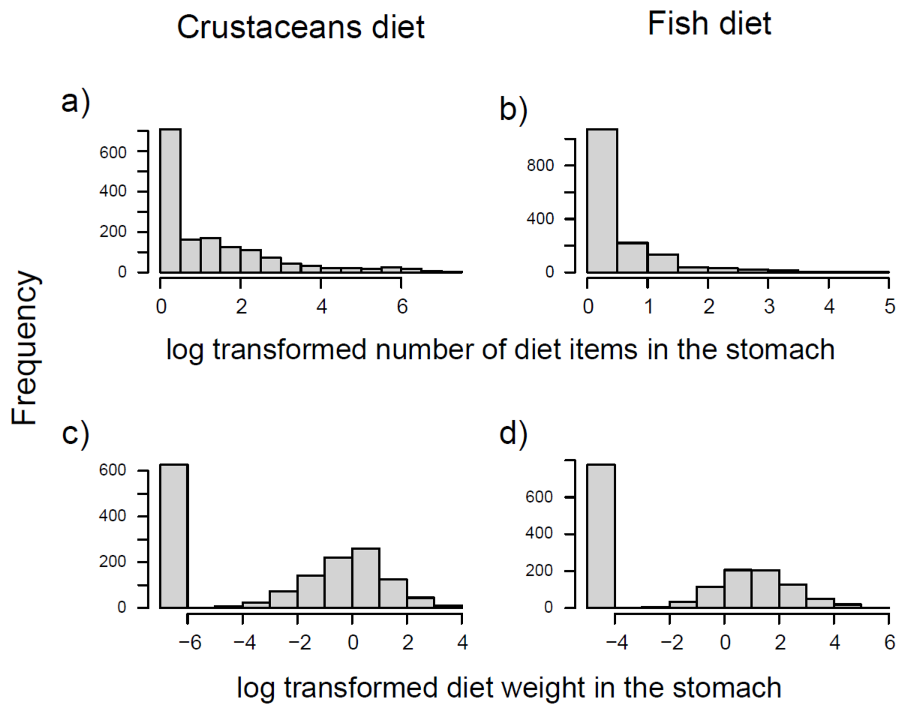


**Fig. S3.** The distribution of log transformed crustaceans and fish diet numbers (a, b) and weights (c, d). Note that one count is added to the number variables, and minimum observed diet weight in the respective datasets (0.002 for crustaceans and 0.01 for fish diet) were added to avoid undefined log (0) value.

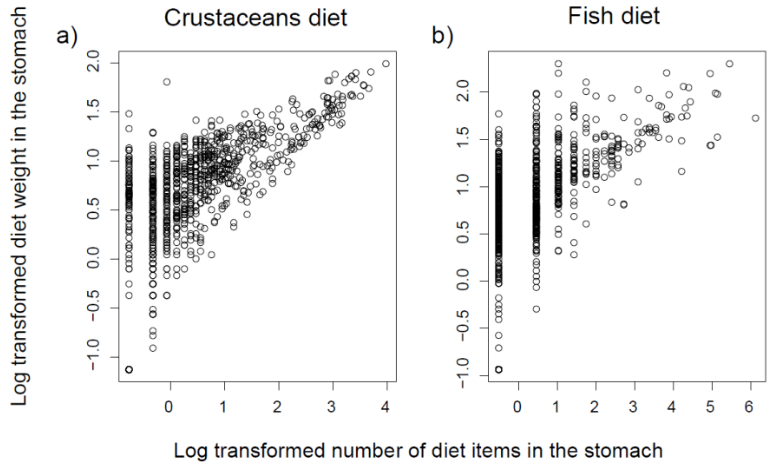


**Fig. S4**. Correlation between log transformed diet number and weight for crustaceans (a) and fish (b) diets. Note that one count is added to the number variables, and minimum observed diet weight in the respective datasets (0.002 for crustaceans and 0.01 for fish diet) were added to avoid undefined log (0) value.

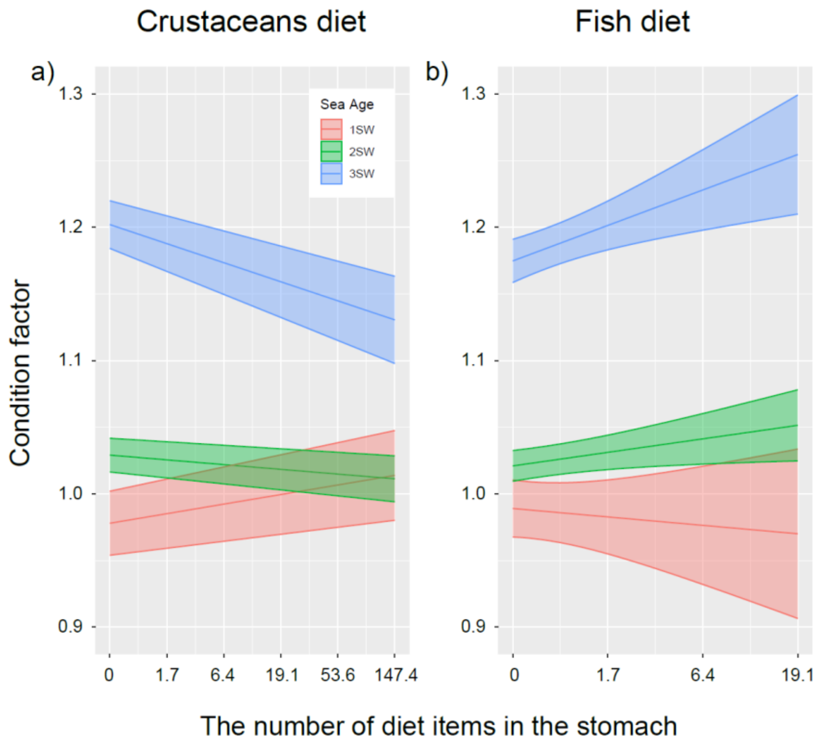


**Fig. S5**. Marginal effects of the number of crustaceans (a) and fish (b) prey items (log scale) on condition factor in Atlantic salmon sub-adults in the marine environment within each age groups. Note that the x-axis indicates diet item numbers on the log scale. Error bars indicate 95% confidence intervals.

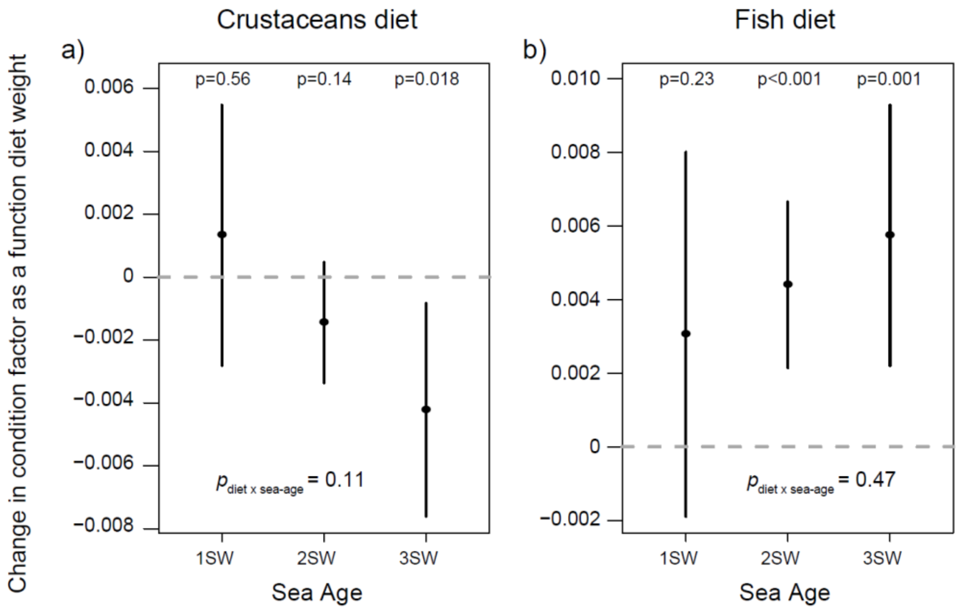


**Fig. S6.** The effect of crustaceans (a) and fish (b) prey weight in the stomach on condition factor in Atlantic salmon sub-adults in the marine environment. Dashed line indicates a zero slope, no relation between diet weight and condition factor. Note that +1 is added to the number variables to avoid undefined log (0) value. Error bars indicate 95% confidence intervals.

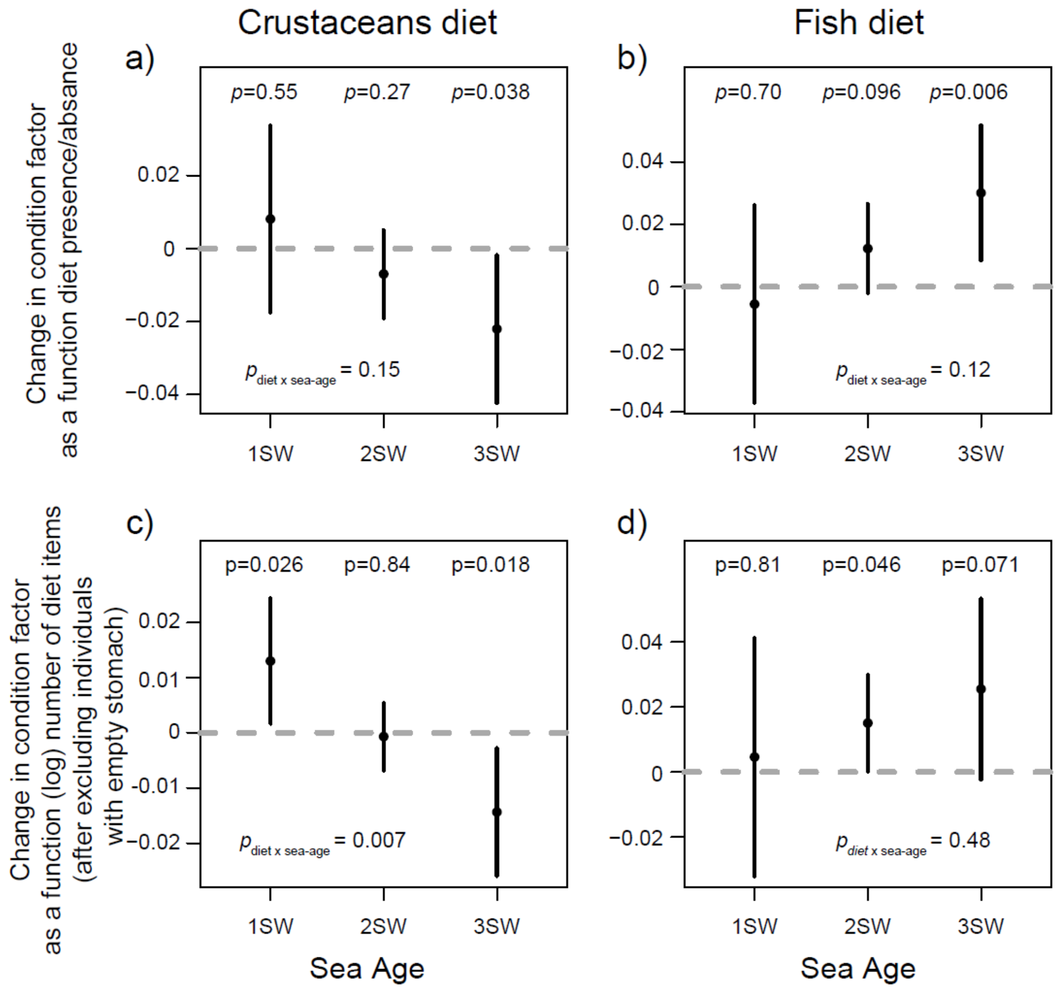


**Fig. S7.** Diet effect on condition when processes parameterized separately; diet presents/absence (Boolean) was modelled as a proxy for foraging frequency (a,b), and when diet present, number of diet items in the stomach (log scale) was modelled as a proxy for foraging efficiency (c,d). . Dashed line indicates a zero slope, no relation between the number of diet items and condition factor. Error bars indicate 95% confidence intervals.

**Table S1.** The descriptive of Atlantic salmon stomachs content in this study (N=1604).

| index | Species | Type | Larger taxonomic groups | $N_{stomach}$ | Weight (g) | $N_{item}$ | Average Weight (g) |
|---|---|---|---|---|---|---|---|
| 1 | *Meganyctiphanes norvegica* | Crustaceans | Euphausiids | 657 | 232.13 | 2043 | 0.114 |
| 2 | *Themisto libellula* | | Hyperiid amphipods | 419 | 201.369 | 2040 | 0.099 |
| 3 | *Themisto compressa* | | Hyperiid amphipods | 378 | 728.947 | 27644 | 0.026 |
| 5 | *Hymenodora glacialis* | | shrimp | 253 | 251.05 | 351 | 0.715 |
| 6 | *Themisto spp* | | Hyperiid amphipods | 217 | 279.908 | 7889 | 0.035 |
| 7 | *Themisto compressa f. bispinosa* | | Hyperiid amphipods | 83 | 43.375 | 2058 | 0.021 |
| 8 | *Themisto compressa f. compressa* | | Hyperiid amphipods | 53 | 23.036 | 867 | 0.027 |
| 9 | *Ephausiidae* | | Euphausiids | 44 | 25.04 | 305 | 0.082 |
| 10 | *Themisto abyssorum* | | Hyperiid amphipods | 27 | 1.86 | 33 | 0.056 |
| 11 | *Eusirus holmi* | | amphipod | 26 | 14.12 | 27 | 0.523 |
| 12 | *Thysanoessa inermis* | | Euphausiids | 17 | 3.83 | 44 | 0.087 |
| 13 | *Sergestes arcticus* | | shrimp | 7 | 5.36 | 7 | 0.766 |
| 14 | *Gammaridea* | | | 3 | 0.93 | 3 | 0.310 |
| 15 | *Pasiphea tarda* | | shrimp | 1 | 0.89 | 1 | 0.890 |
| 16 | *Thysanoessa longicaudata* | | Euphausiids | 1 | 0.004 | 1 | 0.004 |
| 18 | *Maurolicus muelleri* | Fish | Pearlsides | 333 | 983.44 | 888 | 1.107 |
| 19 | *Benthosema glaciale* | | Lanternfishes | 270 | 815.052 | 377 | 2.162 |
| 20 | *Myctophidae* | | Lanternfishes | 123 | 645.17 | 308 | 2.095 |
| 21 | Fry (mostly *Mallotus villosus*) | | | 66 | 111.76 | 424 | 0.264 |
| 22 | *Paralepidae* | | Baracudinas | 37 | 802.27 | 39 | 20.571 |
| 23 | *Myctophum punctatum* | | Lanternfishes | 16 | 44.67 | 23 | 1.942 |
| 24 | *Micromesistius poutassou* | | blue whiting | 12 | 450.39 | 12 | 37.533 |
| 25 | *Notolepis rissoi kroyeri* | | Baracudinas | 9 | 189.64 | 9 | 21.071 |
| 26 | *Clupea harengus* | | Herring | 5 | 360.23 | 5 | 72.046 |
| 27 | *Notoscopelus kroeyeri* | | Lanternfishes | 6 | 52.32 | 7 | 7.474 |
| 28 | *Scomber scombrus* | | Mackerel | 6 | 235.34 | 6 | 39.223 |
| 29 | *Ammodyteidae* | | Sandlance | 3 | 2.71 | 4 | 0.678 |
| 30 | *Gasterosteus aculeatus* | | Three-spined stickleback | 3 | 8.5 | 3 | 2.833 |
| 31 | *Mallotus villosus* | | Capelin | 2 | 40.96 | 3 | 13.653 |
| 32 | *Paralepis coregonoides borealis* | | Baracudinas | 2 | 62.13 | 2 | 31.065 |
| 33 | *Lampanyctus crocodilus* | | Lanternfishes | 1 | 13.52 | 1 | 13.520 |
| 34 | *Lycenchelys spp* | | Eelpouts | 1 | 8 | 1 | 8 |
| 35 | *Gonatidae* | Squid | | 0 | 217.75 | 9* | NA |
| 36 | Remains organic | Remains organic | | 0 | 68.24 | 0* | NA |
| 4 | Crustacea remains | Crustacea remains | | 266 | 274.14 | 13* | NA |
| 17 | Fish remains | Fish | | 550 | 1704.99 | 23* | NA |

* Only a subset of remains were enumerated. When enumerated, fish and squid remains were counted as independent diet data in the fish diet, and crus remains were counted as independent diet entities in the crustaceans group.

**Table S2.** Coefficients of the model with condition as the response variables, and with (log scale) number of items as variables. $Length_{age.std}$ and Sex are scaled. Note that main sea age effect (numeric) is rank-deficient due to confounding effect of the categorized parameterization of the same factor. Note that the table output (obtained from the coefficients summary table) provides z-values, while t-values presented in the text are obtained using emmeans package.

| **Fixed coefficients** | **Estimate** | **SE** | ***z* value** | ***p* value** |
|---|---|---|---|---|
| Intercept | 1.0149 | 0.0251 | 40.39 | < 0.001 |
| Season (winter) | -0.0917 | 0.0113 | -8.12 | < 0.001 |
| Fishing period (93/94) | -0.0182 | 0.0091 | -1.99 | 0.047 |
| Sea Age (SW2) | -0.0036 | 0.0082 | -0.44 | 0.663 |
| Sea Age (SW3) | 0.0463 | 0.0172 | 2.69 | 0.007 |
| $Length_{age.std}$ | NA | NA | NA | NA |
| Sex | 0.0175 | 0.0067 | 2.61 | 0.009 |
| Sea Age (numeric) | -0.0249 | 0.0186 | -1.34 | 0.181 |
| Number of crustaceans diet | 0.0302 | 0.0084 | 3.6 | < 0.001 |
| Number of fish diet | 0.0198 | 0.0098 | 2.02 | 0.043 |
| Sea Age (SW2) : $Length_{age.std}$ | -0.0139 | 0.0059 | -2.36 | 0.018 |
| Sea Age (SW3) : $Length_{age.std}$ | -0.0873 | 0.0189 | -4.62 | < 0.001 |
| Sea Age (SW2) : Sex | -0.0460 | 0.0212 | -2.17 | 0.030 |
| Sea Age (SW3) : Sex | -0.0105 | 0.0033 | -3.21 | 0.001 |
| Sea Age (numeric) : number of crustaceans diet | 0.0174 | 0.0083 | 2.09 | 0.037 |
| Sea Age (numeric) : number of fish diet | 1.0149 | 0.0251 | 40.39 | < 0.001 |

| **Variance components** | **Variance** |
|---|---|
| Regional group | 0.000832 |
| Location | 0.000399 |
| Residuals | 0.012419 |

**Table S3:** Parsimony of null-model structures with different parametrizations of non-focal terms. All combination of fixed effects have been tested, but only models that are ΔAICc < 2 to the best model is displayed in the table. Models were performed both for categorical and continuous parametrization of the sea age effect. Note that best mode with categorical parametrization of sea age was substantially higher than the best model with continuous parametrization (ΔAICc = 67.18).

| Sea Age parameterization | intercept | period | season | $L_{age.std}$ | SeaAge | Sex | season: $L_{age.std}$ | season: SeaAge | season: Sex | $L_{age.std}$: SeaAge | $L_{age.std}$: Sex | SeaAge: Sex | season: $L_{age.std}$: SeaAge | season: $L_{age.std}$: Sex | season: SeaAge: Sex | $L_{age.std}$: SeaAge: Sex | season: $L_{age.std}$: SeaAge: Sex | df | logLik | AICc | delta |
|---|---|---|---|---|---|---|---|---|---|---|---|---|---|---|---|---|---|---|---|---|---|
| categorical | 1.0206 | + | + | -0.0043 | + | 0.0418 | | | | + | -0.0140 | + | | | | | | 15 | 1210.16 | -2390.01 | 0.00 |
| | 1.0252 | + | + | -0.0038 | + | 0.0274 | | | + | + | -0.0168 | + | | | | | | 16 | 1210.99 | -2389.64 | 0.37 |
| | 1.0346 | + | + | -0.0018 | + | 0.0419 | | + | | + | -0.0139 | + | | | | | | 17 | 1211.98 | -2389.57 | 0.44 |
| | 1.0397 | + | + | -0.0012 | + | 0.0277 | | + | + | + | -0.0167 | + | | | | | | 18 | 1212.80 | -2389.16 | 0.85 |
| | 1.0227 | | + | -0.0017 | + | 0.0427 | | + | | + | -0.0140 | + | | | | | | 16 | 1210.40 | -2388.46 | 1.55 |
| | 1.0224 | + | + | -0.0016 | + | 0.0428 | + | | | + | -0.0140 | + | | | | | | 16 | 1210.37 | -2388.4 | 1.60 |
| | 1.0273 | + | + | -0.0008 | + | 0.0281 | + | | + | + | -0.0170 | + | | | | | | 17 | 1211.25 | -2388.12 | 1.89 |
| continuous | 0.9571 | | + | 0.0219 | 0.0662 | 0.0117 | | + | + | -0.0001 | -0.0480 | -0.0241 | | | | 0.0150 | | 14 | 1175.55 | -2322.83 | 0.00 |
| | 0.9582 | | + | 0.0062 | 0.0657 | 0.0109 | | + | + | 0.0075 | -0.0167 | -0.0231 | | | | | | 13 | 1174.37 | -2322.51 | 0.32 |
| | 0.9725 | | + | 0.0217 | 0.0585 | 0.0044 | | + | + | | -0.0164 | -0.0193 | | | | | | 12 | 1173.31 | -2322.42 | 0.41 |
| | 0.9532 | | + | 0.0203 | 0.0646 | 0.0257 | | + | | 0.0000 | -0.0423 | -0.0205 | | | | 0.0142 | | 13 | 1174.05 | -2321.87 | 0.96 |
| | 0.9683 | | + | 0.0205 | 0.0573 | 0.0177 | | + | | | -0.0127 | -0.0161 | | | | | | 11 | 1172.02 | -2321.87 | 0.96 |
| | 0.9544 | | + | 0.0055 | 0.0642 | 0.0244 | | + | | 0.0072 | -0.0129 | -0.0197 | | | | | | 12 | 1173.01 | -2321.81 | 1.01 |
| | 0.9635 | + | + | 0.0223 | 0.0665 | 0.0114 | | + | + | -0.0004 | -0.0481 | -0.0240 | | | | 0.0151 | | 15 | 1175.91 | -2321.51 | 1.31 |
| | 0.9827 | | + | 0.0201 | 0.0502 | -0.0164 | | + | | | -0.0132 | | | | | | | 10 | 1170.75 | -2321.36 | 1.47 |
| | 0.9790 | + | + | 0.0216 | 0.0590 | 0.0043 | | + | + | | -0.0163 | -0.0193 | | | | | | 13 | 1173.72 | -2321.21 | 1.62 |
| | 0.9646 | + | + | 0.0065 | 0.0659 | 0.0107 | | + | + | 0.0073 | -0.0166 | -0.0230 | | | | | | 14 | 1174.72 | -2321.18 | 1.65 |
| | 0.9691 | | + | 0.0229 | 0.0600 | -0.0173 | | + | + | -0.0006 | -0.0520 | -0.0089 | | | + | 0.0170 | | 15 | 1175.72 | -2321.14 | 1.69 |
| | 0.9881 | | + | 0.0210 | 0.0500 | -0.0320 | | + | + | | -0.0161 | | | | | | | 11 | 1171.55 | -2320.94 | 1.89 |

**Table S4:** Model parsimony between different parameterization crustaceans, modelled either as a whole group (crustaceans) or hyperiid amphipods and euphausiids separately. Note that the model structure is based on the best model presented Table 1.

| **Contrasting diet parameter(s) between models** | **Model structure** | **LL** | **df** | **AICc** | **dAICc** |
|---|---|---|---|---|---|
| Crustacean | Null model parameters + $SeaAge_{continious}$ x ($Diet_{crus}$ + $Diet_{fish}$) | 1221.464 | 19 | -2404.45 | 2.53 |
| Hyperiid amphipods | Null model parameters + $SeaAge_{continious}$ x ($Diet_{Hyperiid.amphipods}$ + $Diet_{fish}$) | 1222.731 | 19 | -2406.98 | 0 |
| Euphausiids | Null model parameters + $SeaAge_{continious}$ x ($Diet_{amphipods}$ + $Diet_{fish}$) | 1215.232 | 19 | -2391.98 | 15.00 |
| Hyperiid amphipods and Euphausiids | Null model parameters + $SeaAge_{continious}$ x ($Diet_{Hyperiid.amphipods}$ + $Diet_{amphipods}$ + $Diet_{fish}$) | 1223.028 | 21 | -2403.47 | 3.51 |
